# Human Papillomavirus Type 16 L1/L2 VLP Experimental Internalization by Human Peripheral Blood Leukocytes

**DOI:** 10.1101/299214

**Authors:** Aurora Marques Cianciarullo, Vivian Szulczewski, Erica Akemi Kavati, Tania Matiko Hosoda, Elizabeth Leão, Primavera Borelli, Enrique Boccardo, Martin Müller, Balasubramanyam Karanam, Willy Beçak

## Abstract

Human papillomavirus (HPV) accounts for hundreds of thousands of new cases of cervical cancer yearly, and half of these women die of this neoplasia. This study investigates the possibility of HPV16 L1/L2VLP to be internalized by human peripheral blood leukocytes in *ex vivo* assays. We have developed a leukocyte separation method from heparinized blood samples aiming cellular integrity and viability. We have expressed humanized L1 and L2 viral capsid proteins in HEK293T epithelial human cells, transiently transfecting them with vectors encoding humanized HPV16 *L1* and *L2* genes. Recombinant L1/L2 capsid proteins and structured virus-like particles interacted with human peripheral blood mononuclear cells, lymphocytes and monocytes, and were internalized through a pathway involving CD71 transferrin receptors. This was observed, at a percentile of about 54% T- CD4, 47% T-CD8, 48% B-CD20, and 23% for monocytes-CD14. The group of polymorph nuclear cells: neutrophils-eosinophils-basophils group did not internalize any VLPs. Blockage assays with biochemical inhibitors of distinct pathways, like chlorpromazine, rCTB, filipin, nystatin, liquemin, and sodium azide also evidentiated the occurrence of virus-like particles indiscriminate entrance via membrane receptor on mononuclear cells. This study shows that HPV16 L1/L2 VLPs can interact with the plasma membrane surface and successfully enter lymphocytes without requiring a specific receptor.

**Legend of the Graphical Abstract:** Graphical abstract showing *ex vivo* and *in vitro* internalization between VLPs and host cells.

After leukocytes separation from human whole blood, it was performed the identification of human peripheral blood leukocytes in ex vivo interactions with VLPs. The graph shows that of the cells that interacted with VLPs, 52% corresponded to lymphocytes T-CD4, 47% lymphocytes T-CD8, 48% lymphocytes B-CD20, and only 23% of the monocytes CD14 interacted with these particles. However, monocytes apparently internalized larger amounts of particles when compared to lymphocytes.

It is probable that in some T lymphocytes the amount of internalized particles has been imperceptible to the confocal microscope, since the VLPs produced in this research are around 50 nm in diameter. These results lead to two important implications. First, the interaction of VLPs with lymphocytes may result in the activation of these cells and, consequently, increase the population of these circulating cells, this being crucial in the induction of specific immune response.

In the second implication, these lymphocytes would internalize small amounts of virus, insufficient to activate the immune system. Here it is important to note that lymphocytes are cells capable of dividing and it is estimated that the half-life of these inactive cells in humans is of some years. In addition, as it is known, inactive lymphocytes continually re-circulate through the bloodstream and lymphatic vessels.

The percentage of cells that interacted with the HPV16 L1/L2 VLPs was calculated by the number of cells recognized by the anti-CD antibodies, which internalized these particles. The result corresponds to the analysis in duplicates, being representative of at least four tests.

All images are original and cells were processed by Cianciarullo AM et al., at the Butantan Institute, Sao Paulo – SP, Brazil.

Electron micrographs of human leukocytes, HEK293T and HPV16 L1/L2 VLPs were obtained in a Zeiss EM109 transmission electron microscope. The blue color of the VLPs and colored leukocytes were virtually attributed. Leukocyte and HEK293T present filamentous actin (red) and HPV16 L1/L2 VLPs (green), by fluorescence in a Confocal Zeiss LSM 510 Meta Microscope

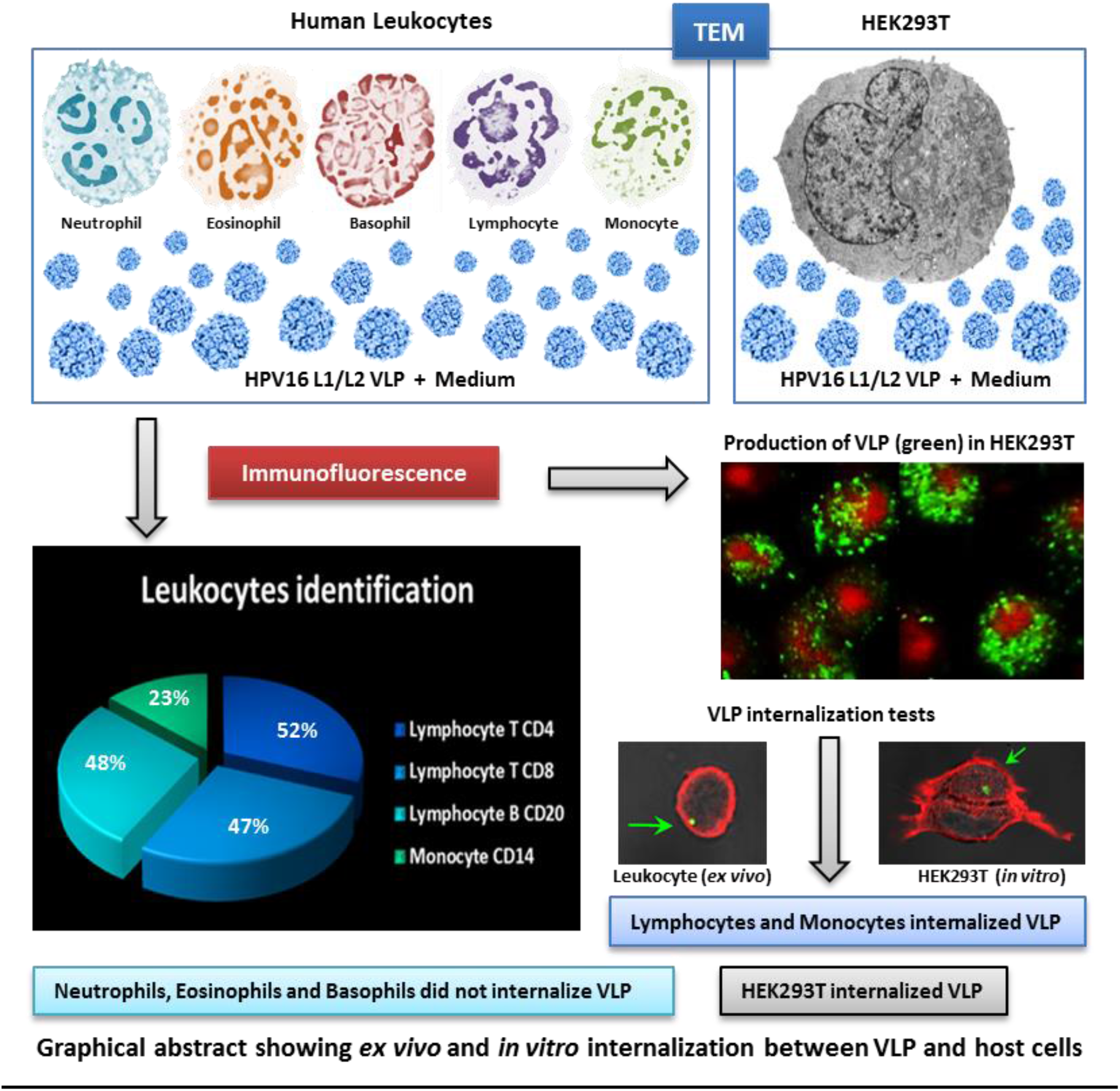

## INTRODUCTION

In 1974, zur Hausen researches pointed to the HPV as the major etiological agent in cervical cancer and, later, its DNA was detected in tumors found in other anogenital regions (1–3). Today, there are more than 200 different types of well characterized HPVs, of which approximately 40 types infect the genital tract. The most frequently found are types 6 and 11 that induce benignant skin warts formation, and types 16 and 18 which are associated to cancer development. There are at least 12 HPV types that according to the Agency for Research on Cancer (IARC) are considered oncogenic to humans (4, Papillomavirus Episteme (PaVE); http://pave.niaid.nih.gov/). Among them, HPV16 and HPV18 are those responsible for approximately 70% of all cervical cancers (4, 5). In 2025, the projected global estimate of cervical cancer is expected to rise to 720,415 new cases per year, and it is expected that half of these women will die of this neoplasia (5). More recently, novel HPV lineages and sub-lineages were described, found in HIV/HPV-co-infected pregnant women, with two high-risk types (6). However, the difficulty to obtain enough viable wild types or recombinant HPV particles has limited researches to distinct aspects of virus biology (7). All viruses enter host cells to survive, replicate and evade the immune system. So far, the entry of HPV and its traffic through the host cells are still not completely elucidate (8). Several studies describe this process like a complex set of interactions among different pathways, receptors, co-receptors and co-factors. In keratinocytes, HPV seems to be internalized via clathrin-dependent endocytic mechanisms, but it might use alternative uptake pathways to enter cells, such as a caveolae-dependent route, among others depending on viral type (7–10).

Currently, HPV is recognized as one of the main causes of infection-related cancer worldwide, as well as the causal factor of other diseases. Infection with high-risk HPV types is the etiological cause of cervical cancer and is strongly associated with a significate fraction of penile, vulvar, vaginal, anal and oropharyngeal cancers (11, 12). HPVs are responsible for approximately 88% of anal cancer and 95% of anal intraepithelial neoplasia grades 2/3 lesions, and 40%-50% of penis and vulvar cancers (11). In oropharyngeal cancers, HPV DNA was detected in 35%-50% cases (11). In all HPV-positive non-cervical cancers, HPV16 is the most common HPV type detected, followed by HPV types 18, 31, 33 and 45. Among the non-cancerous HPV-associated conditions, genital warts and recurrent respiratory papillomatosis are surely linked to HPV6 and 11 (10). In addition, it was demonstrated recently that HPV16 virus-like particles (VLPs) L1/L2 interact with hematopoietic precursor stem cells, present in the amniotic fluid from healthy pregnant women (13). Whereas new pathologies are increasingly being associated to the HPVs; the responsibility and costs of HPV-associated diseases and cancer remain an important public health issue in all countries, regardless of their economic developmental level (14). Due to the continuous worldwide propagation of HPV it is necessary to investigate the possibility of HPV internalization by different human cell types. In the present study, we addressed the capacity of HPV16 L1/L2 VLP entrance/uptake of – in peripheral blood leukocytes *ex vivo*.

## MATERIALS AND METHODS

### Production of HPV16 L1/L2 VLPs

The production of VLPs was carried out throughout the recombinant protein expression HPV16 L1 and L2 in epithelial human cells of the HEK293T lineage, cultured as previously described (15). Transfection of these cells occur in a transient way using pUF3L1h and pUF3L2h vectors, which are regulated by the human cytomegalovirus promoter containing complete sequences of genes *L1* and *L2*, a method already described in detail (15, 16). In this study, besides being used to produce VLPs containing both capsid proteins, HEK293T cell line was also used as a control factor on internalization assays with human peripheral blood leukocytes. All assays performed in this study were analyzed in duplicate, being representative of at least four independent tests.

### Blood collection

Blood samples from 10 healthy female volunteers, ages ranging from 35 to 55, were requested based on epidemiological data associated with genital HPV infection (17). The screening of the volunteers was based on recent blood, Pap smears and colposcopy tests and on data filled out by the candidate on a written informed consent form and on a questionnaire, for the purpose of laboratory research. Volunteers who did not present Pap and colposcopy results within reference values were dismissed. Blood samples were collected in sterile tubes containing heparin, for internalization assays with leukocytes, and EDTA for blocking assays, and were processed quickly, within a maximum of 2-hours after collection. During this interval, they were stored at 4°C.

### Leukocytes separation

Whole blood was collected in heparin at a concentration of 0.1 mg/ml, centrifuged at 1,000 rpm for 7 minutes at 5°C in a Sorvall^®^ RT 6000 Refrigerated Centrifuge (Du Pont, Wilmington-DE, USA) with a horizontal angle rotor. The plasma supernatant containing platelets and leukocytes was gently removed and transferred with the aid of a Pasteur pipette to a new tube. An aliquot was collected for total and differential counts, performed in a Neubauer chamber, diluted (ratio 1:1) in Trypan blue and smears were stained by May-Grünwald-Giemsa in order to determine their composition. Leukocytes sedimented were centrifuged again at 1,200 rpm, for 3 minutes, at 5°C, to remove platelets. Leukocytes precipitates were resuspended in 0.85% saline solution for subsequent tests (18). The Neubauer chamber counts and smears containing distinct cell types were analyzed by light microscopy, with a Leica DMIL I microscope (Leica Microsystems GmbH, Vienna, AUT).

### Internalization assays for HEK293T cells and HPV16 L1/L2 VLPs

HEK293T cells (2×10^4^cells/ml) were plated and maintained under growth conditions, washed with PBS and incubated with 120 µg of VLPs in DMEM without FBS, for 4-hours at 37°C and 5% CO_2_. After this, cells were washed twice with PBS for 3 minutes each in order to remove non-internalized particles. Cells were then fixed with 2% PFA (Paraformaldehyde, Sigma-Aldrich) in PBS for 1-hour, at 4°C, and washed three times with PBS, 5 minutes each. Plates were kept at 4°C until immunofluorescence assays. Controls were performed in the absence of VLPs and/or the denaturation thereof, by heating at 100°C for 10 minutes (19, adapted).

### Internalization assays for human leukocytes and HPV16 L1/L2 VLPs

Leukocytes (2×10^4^ cells/ml) of healthy volunteers were incubated with 120 µg of VLPs produced in this study and RPMI without FBS for 4-hours at 37°C and 5% CO_2_, under gentle agitation. Cells were centrifuged at 1,000 rpm for 7 minutes and cell pellets washed three times with PBS, 5 minutes each. Then, cells were fixed with 2% PFA in PBS for 1-hour, at 4°C. After this step, the centrifugation and PBS washing processes were repeated. Samples were kept at 4°C until immunofluorescence assays. Controls were the same as described above for HEK293T cells.

### Internalization assays for leukocytes, transferrin and HPV VLPs

The same protocol described above was employed, with minor changes described below. Leukocytes were incubated with 120 µg of VLPs in separate samples, as follows: the HPV16 L1/L2 VLPs produced in this study, and L1 VLPs of HPV6, 11, 16 and 18 from Gardasil^®^ vaccine, used as control, kindly provided by Merck Sharp & Dohme. RPMI without FBS, together with transferrin (Tf) conjugated to fluorochrome TexasRed^®^ (Molecular Probes™), were added (ratio 1:60, Tf:RPMI). Samples were kept for 15, 45, 60 and 120 minutes in an incubator at 37°C and 5% CO_2_, under gentle agitation. The other procedures remained unchanged.

### Blocking assays for membrane receptors

Leukocytes of healthy donors were counted and incubated in RPMI medium overnight over coverslips containing poly-L-lysine (Sigma-Aldrich), at 37°C and 5% CO_2_. Then, cells were washed and a fresh medium was placed together with specific biochemical inhibitors of ligand uptake (Table 1), used isolated or associated, and incubated for 2 hours (19). After incubation, cells were washed and 120 µg of HPV16 L1/L2 VLPs (20) were added and incubated with medium for 4 hours. Cells were washed again and fixed in 2% PFA solution. Immunofluorescence assays were carried out to detect the VLP-PBMC interaction by confocal microscopy, using specific antibodies to recognize L1 and L2 proteins. Z-axis 3D images were obtained to confirm the presence of VLPs within cells.

**Table 1:** List of primary antibodies used in immunocytochemistry reactions.

| Antibody | Dilution | Description | Supplier |
| --- | --- | --- | --- |
| **Anti-L1**<br>**Camvir-1** | 1:500 | anti-mouse IgG<sub>2a</sub>, specific for HPV16 | BD Biosciences |
| **Anti-L1**<br>**conformational** | 1:100 | anti-mouse IgG<sub>2a</sub>, specific for HPV16 | Biodesign |
| **Anti-L2** | 1:100 | anti-rabbit IgG | Roden, R.<br>MD/USA |
| **Transferrin**<br>**conjugated** | 1:60 | TexasRed<sup>®</sup> (Ex. 595 nm, Em. 615 nm) | Molecular<br>Probes<sup>™</sup> |
| **Anti-Tf** | 1:750 | anti- rabbit IgG | Dako |
| **Anti-Ferritin** | 1:750 | anti- rabbit IgG | Dako |
| **Anti-CD71** | 1:750 | anti-mouse IgG<sub>1</sub>; Tf receptor | Dako |
| **Anti-CD04** | 1:20 | anti-mouse IgG<sub>2a</sub>; recognizes T lymphocytes “helper” | CALTAG<sup>™</sup><br>Laboratories |
| **Anti-CD08** | 1:20 | anti-mouse IgG<sub>2a</sub>; recognizes cytotoxic T lymphocytes | CALTAG<sup>™</sup><br>Laboratories |
| **Anti-CD14** | 1:20 | anti-mouse IgG<sub>2a</sub>; recognizes monocytes | CALTAG<sup>™</sup><br>Laboratories |
| **Anti-CD20** | 1:20 | anti-mouse IgG<sub>3</sub>; recognizes B lymphocytes | CALTAG<sup>™</sup><br>Laboratories |

### Immunofluorescence assays for internalization analysis

Leukocytes were washed and fixed as already described. Samples were incubated in PBS containing 1% BSA for 5 minutes under gentle agitation. Later, cells were incubated with the primary antibodies (Table 1); diluted in PBS containing 0.01% Tween^®^ 20 and 0.5% BSA pH 8 for 2 hours, under gentle agitation, at room temperature. They were then washed three times with PBS for 10 minutes each, followed by incubation with the corresponding secondary antibodies (Table 2) conjugated with fluorochromes and diluted in PBS containing 0.01% Tween^®^ 20 and 1.5% BSA for 1 hour, under light stirring at room temperature. Once more, they were washed three times with PBS for 10 minutes each. Samples of leukocytes with transferrin and VLPs were labelled with Phalloidin conjugated AlexaFluor^®^ 594, incubated for 20 minutes. Immediately after that, they were rinsed twice with PBS for 10 minutes each. Aliquots of cell suspension from both items assayed were adhered over silanized slides, and mounted with 5 µl Mowiol^®^ and coverslips. Samples were kept at 4°C until CLSM analysis, using Confocal Laser Scanning Microscope Zeiss 510 Meta of the Butantan Institute (FAPESP Process No. 2000/11624-5; Carl Zeiss GmbH, Jena, DEU).

**Table 2:**
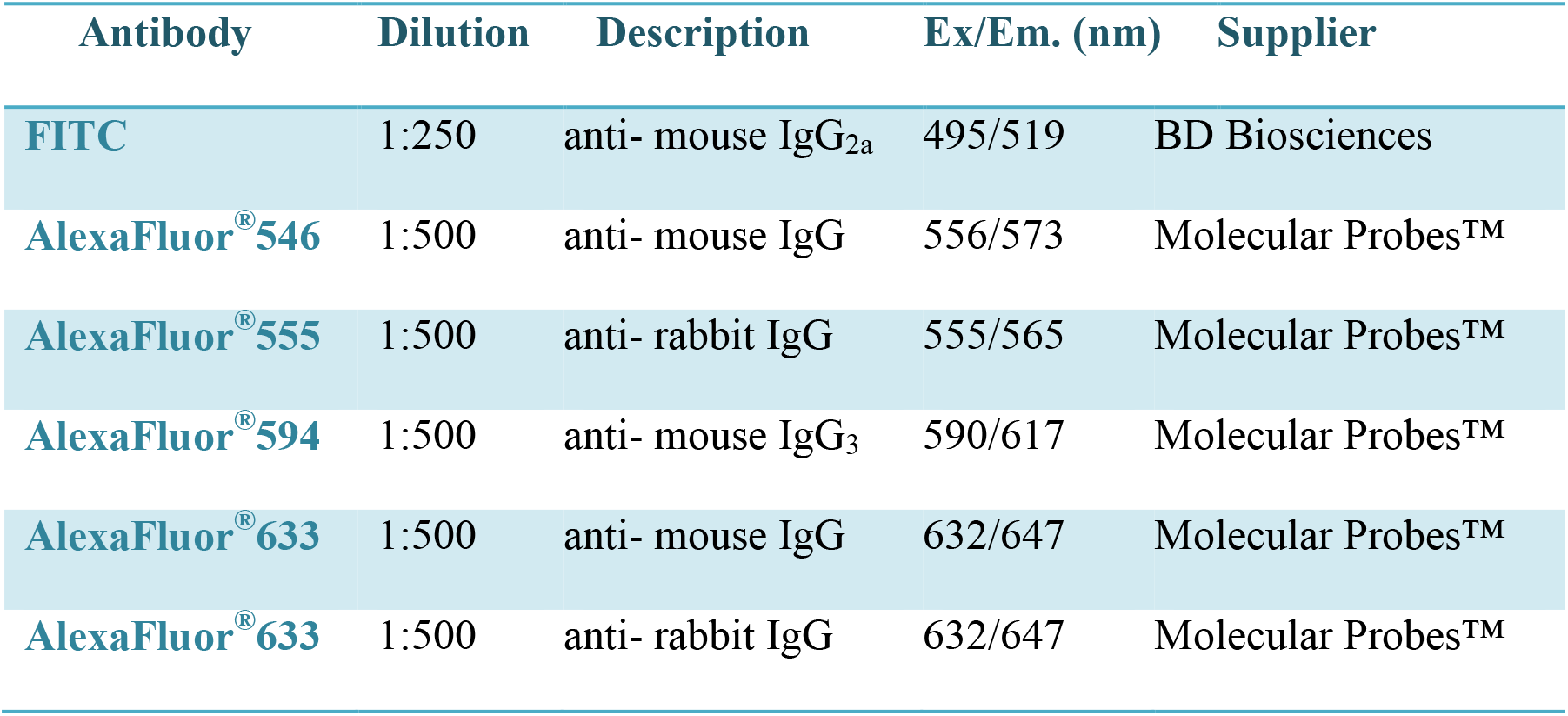
List of secondary antibodies used in immunocytochemistry reactions.

| Antibody | Dilution | Description | Ex/Em. (nm) | Supplier |
| --- | --- | --- | --- | --- |
| **FITC** | 1:250 | anti- mouse IgG<sub>2a</sub> | 495/519 | BD Biosciences |
| **AlexaFluor<sup>®</sup> 546** | 1:500 | anti- mouse IgG | 556/573 | Molecular Probes<sup>™</sup> |
| **AlexaFluor<sup>®</sup> 555** | 1:500 | anti- rabbit IgG | 555/565 | Molecular Probes<sup>™</sup> |
| **AlexaFluor<sup>®</sup> 594** | 1:500 | anti- mouse IgG<sub>3</sub> | 590/617 | Molecular Probes<sup>™</sup> |
| **AlexaFluor<sup>®</sup> 633** | 1:500 | anti- mouse IgG | 632/647 | Molecular Probes<sup>™</sup> |
| **AlexaFluor<sup>®</sup> 633** | 1:500 | anti- rabbit IgG | 632/647 | Molecular Probes<sup>™</sup> |

## RESULTS

### Leukocytes identification

After isolation of leukocytes from healthy donors, control smears stained by May-Grünwald-Giemsa method showed cell morphology preservation (18), presenting a positive correlation with the control blood smears. Counts indicated approximately 98% lymphocytes; 0.7% monocytes; 1.5% polymorph nuclear cells; 0.1% red blood cells and 0.1% platelets. The ultrastructure was well preserved for all cell types analyzed, with well-defined and intact membranes (18). The efficiency and speed of the method for obtaining leukocytes, preserving morphology and cell viability was confirmed.

### Analysis of HPV16 L1/L2 VLPs internalization by HEK293T cells

HPV16 L1/L2 VLPs internalization by HEK293T cells was analyzed through immunofluorescence. We emphasize that the genome of this cell line does not contain any HPV DNA sequences, but it contains Adenovirus DNA and SV40 T-Ag. After 4-hours kept in contact at 37°C, cells were washed in order to remove any particles or proteins not internalized by HEK293T cells. Then, cells were fixed and stained with the primary antibody Camvir-1 (BD Biosciences – Table 1) for L1 proteins and with L1 conformation-specific anti-VLP antiserum (Biodesign – Table 1). It was therefore possible to observe the internalization of VLPs in some cells. In these cells VLPs were found in the cytoplasm, near the core region, stained with anti-VLP. These results were confirmed by the overlap of images, and by the most thoroughly detailed internalization display on the Z-axis scanning sections. The morphological evaluations show that probably the VLPs internalization occurs simultaneously with a large number of particles, similar to the formation of endocytic vesicle structures. These observations suggest that after 4 hours, structured VLPs, as well as non-structured VLPs, such as pentameric and monomeric forms of L1 stained with Camvir-1 (Table 1), were internalized in epithelial cells from the human kidney (HEK293T), across the cell membrane. The negative controls showed no immunostaining for L1 and VLPs and no changes in cell morphology were detected.

### Analysis of HPV16 L1/L2 VLPs internalization by human leukocytes

L1/L2 particles were added to human leukocytes, for 4-hours at 37°C, and prepared for indirect CLSM immunofluorescence analysis, in order to investigate the possibility of internalization between the leukocytes and the HPV16 L1/L2 VLPs. Results indicate an internalization of VLPs with peripheral blood mononuclear cells (PBMC), T and B lymphocytes and monocytes, from healthy women volunteers (Fig. 1). Structured HPV16 L1/L2 VLPs also interacted with and were internalized by leukocytes (Figs. 2 and 3). After 4-hours of interaction, in most leukocytes examined, these particles were found in the cell cytoplasm, as was the case with HEK293T. These results were confirmed by the images overlap (Figs. 1 A, B and D; Figs. 2 and 3), and the detailed internalization, as shown on the Z-axis scanning sections (Fig. 2 C). Fig. 1 C (green arrow) shows VLPs across the cell membrane. The Z-axis sweep cuts were able to demonstrate the internalization of the VLP with a larger number of particles similar to the formation of endocytic vesicles structures (Fig. 2 A, white arrow), like those found in HEK293T cells, a phenomenon endorsed by morphological evaluations. In Fig. 3, the interaction of leukocytes with the VLPs is illustrated, showing the colocalization of HPV16 L1/L2 proteins recognized by anti-L1 (green) and anti-L2 (red), respectively. This colocalization was expected in VLPs that are composed of these two proteins, as it might be seen from the images overlap (Fig. 3). Fig. 3 B shows the internalization of VLPs by leukocytes in a significant amount when compared to other internalizations. At least 15 fields were analyzed per experiment, containing from 4 to 10 cells per field. The presence of structures similar to vacuoles can also be observed (Fig. 3 B, blue arrow). Control assays showed no immunostaining, neither with the antibodies used to detect L1 and L2 separately, nor with structured HPV16 L1/L2 VLPs.

**Figure 1:**
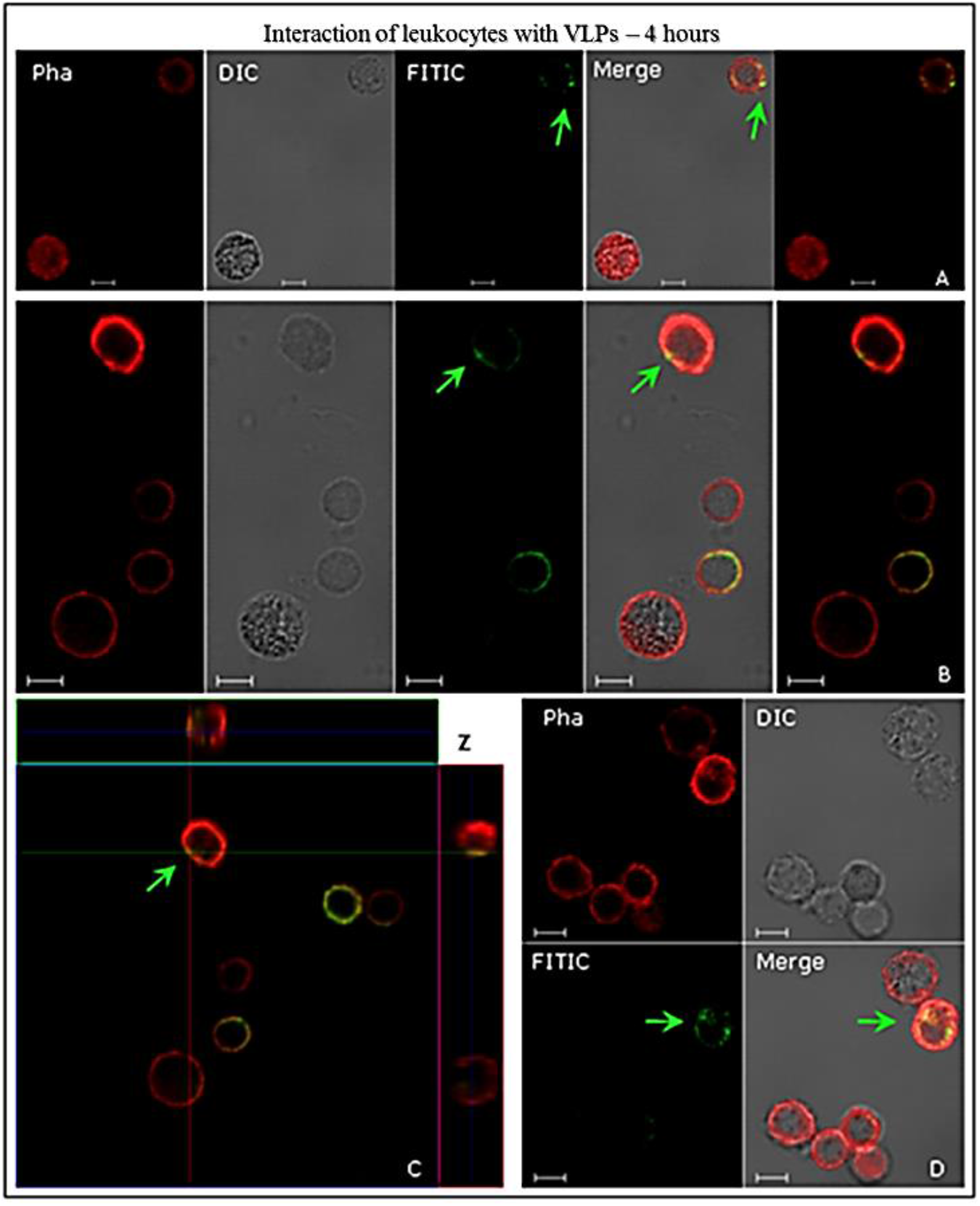
Interactions of human leukocytes with VLPs of HPV16, detected by indirect immunofluorescence. The leukocyte cells were incubated with the VLPs for 4 h. Following fixation, they were immunostained with anti-HPV16 L1 antibody and then stained with a secondary antibody conjugated to FITC (green). In all assays, AlexaFluor^®^ 594-conjugated Phalloidin (red) detected the actin cytoskeleton. The internalization of VLPs (green arrows) is shown in image C, through the sweep cuts on the Z-axis (C) 9.50 µm thick sections and each section to 1.60 µm. CLSM Zeiss LSM 510 Meta. Magnification: Objective C-Apochromatic 63xs /1.4 oil. Bar = 5 µm.

**Figure 2:**
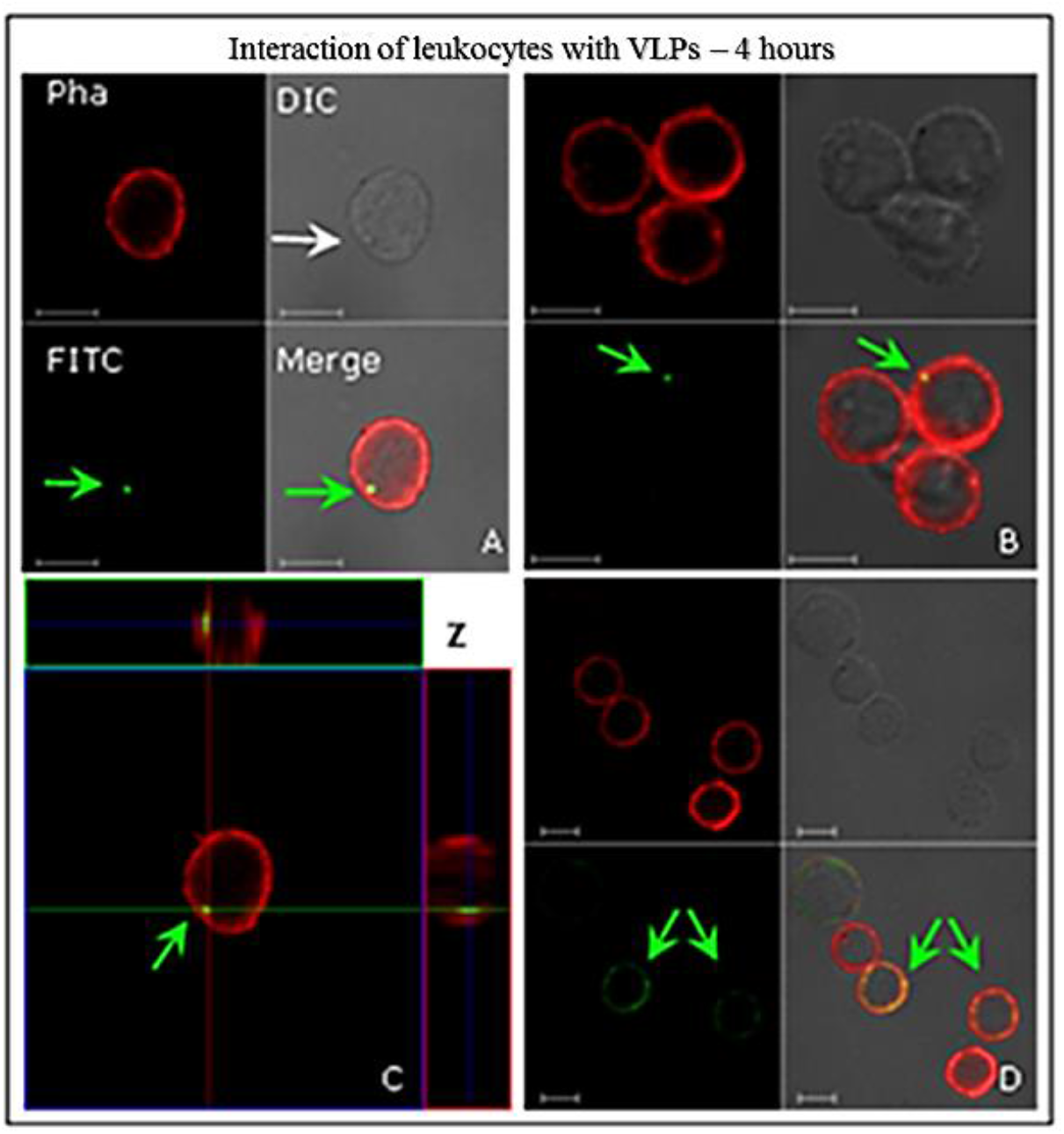
Interactions of human leukocytes with VLPs of HPV16, detected by indirect immunofluorescence. The leukocyte cells were incubated with the VLPs for 4 h. Following fixation, they were immunostained with the conformational anti-HPV16 VLPs and stained with a secondary antibody conjugated to FITC (green). AlexaFluor^®^ 594-conjugated Phalloidin (red) detected the actin cytoskeleton. The internalization of VLPs (green arrows) is shown in image C, through Z-axis sweep cuts (C) 7.80 µm thick sections and each section to 1.30 µm. In (A) we see a structure similar to an endocytic vesicle (white arrow). CLSM Zeiss LSM 510 Meta. Magnification: Objective C-Apochromatic 63xs /1.4 oil. Bar = 5 µm.

**Figure 3:**
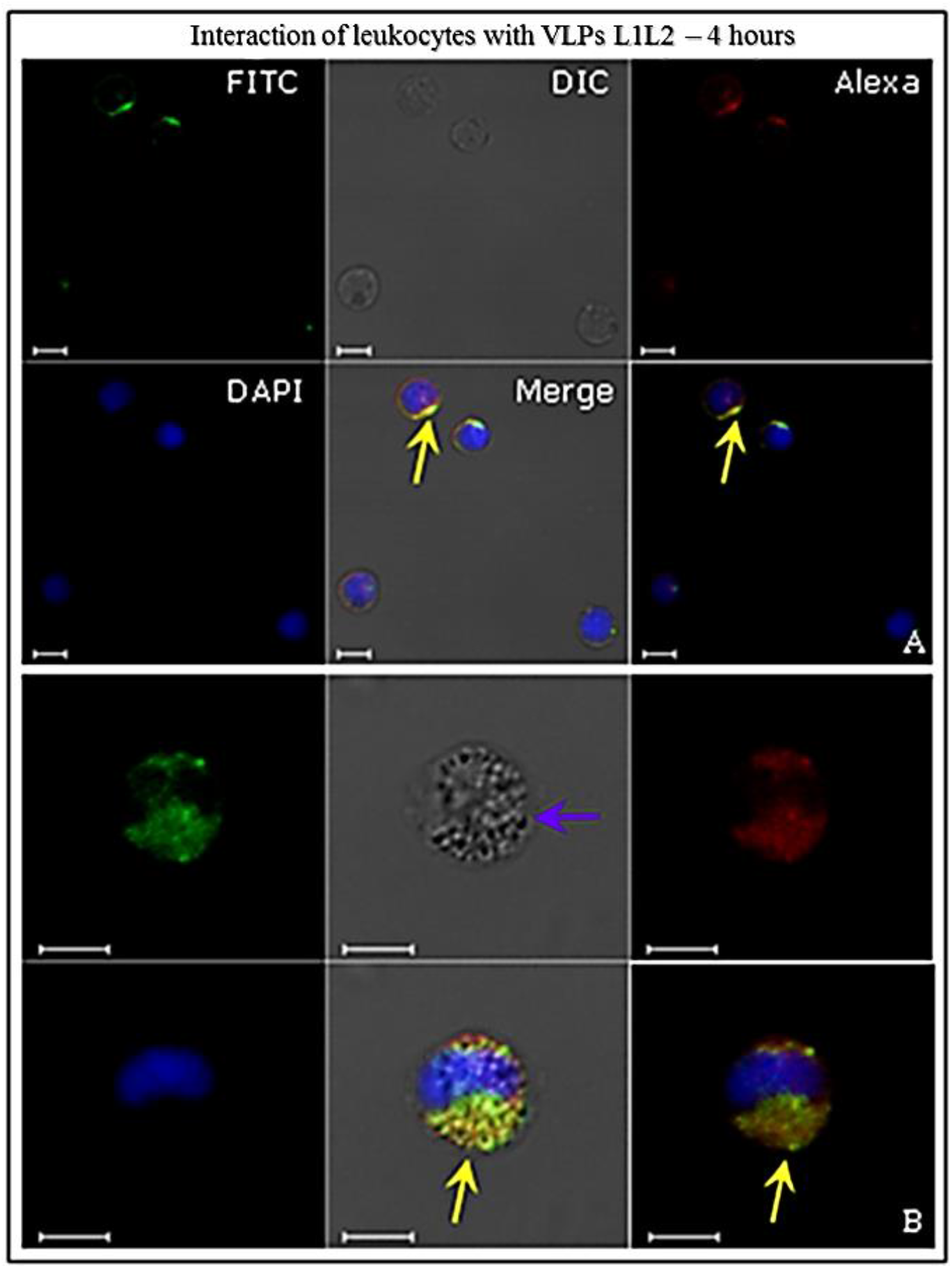
Interactions of human leukocytes with VLPs of HPV16, detected by indirect immunofluorescence. After fixation, cells were immunostained with anti-HPV16 L1 antibody and then stained with a secondary antibody conjugated to FITC (green). Then immunostained with anti-HPV16 L2 antibody and revealed by AlexaFluor^®^555 secondary antibody (red). The internalization of VLPs (yellow arrows) is shown in confocal images (merge). In (B) structures similar to vacuoles (blue arrow) might be observed. CLSM Zeiss LSM 510 Meta. Magnification: Objective C-Apochromatic 63xs /1.4 oil. Bar = 5 µm.

### Identification of VLPs internalization by human PBMC

These cells were treated with antibodies in order to recognize specific cell membrane receptors for each type (Table 1 and Table 2), and analyzed by indirect immunofluorescence using CLSM to identify PBMC interactions with HPV16 VLPs, after 4-hours incubation at 37°C. HPV16 VLPs interacted with *ex vivo* PBMC (Figs. 4–6). These results were confirmed by the images overlap (Figs. 4 and 5) and displayed detailed internalization on the Z-axis sections (Fig. 6). T and B lymphocytes showed greater competence to internalize VLPs, at around 47%-52% of cells (Table 3). Through morphological assessments, T-lymphocytes that internalized VLPs and were recognized by anti-CD8 showed endocytic vesicles-like structures (Fig. 4 B, white arrow). Only 23% of monocytes that were identified by anti-CD14 antibody interacted with the VLPs (Table 3).

**Figure 4:**
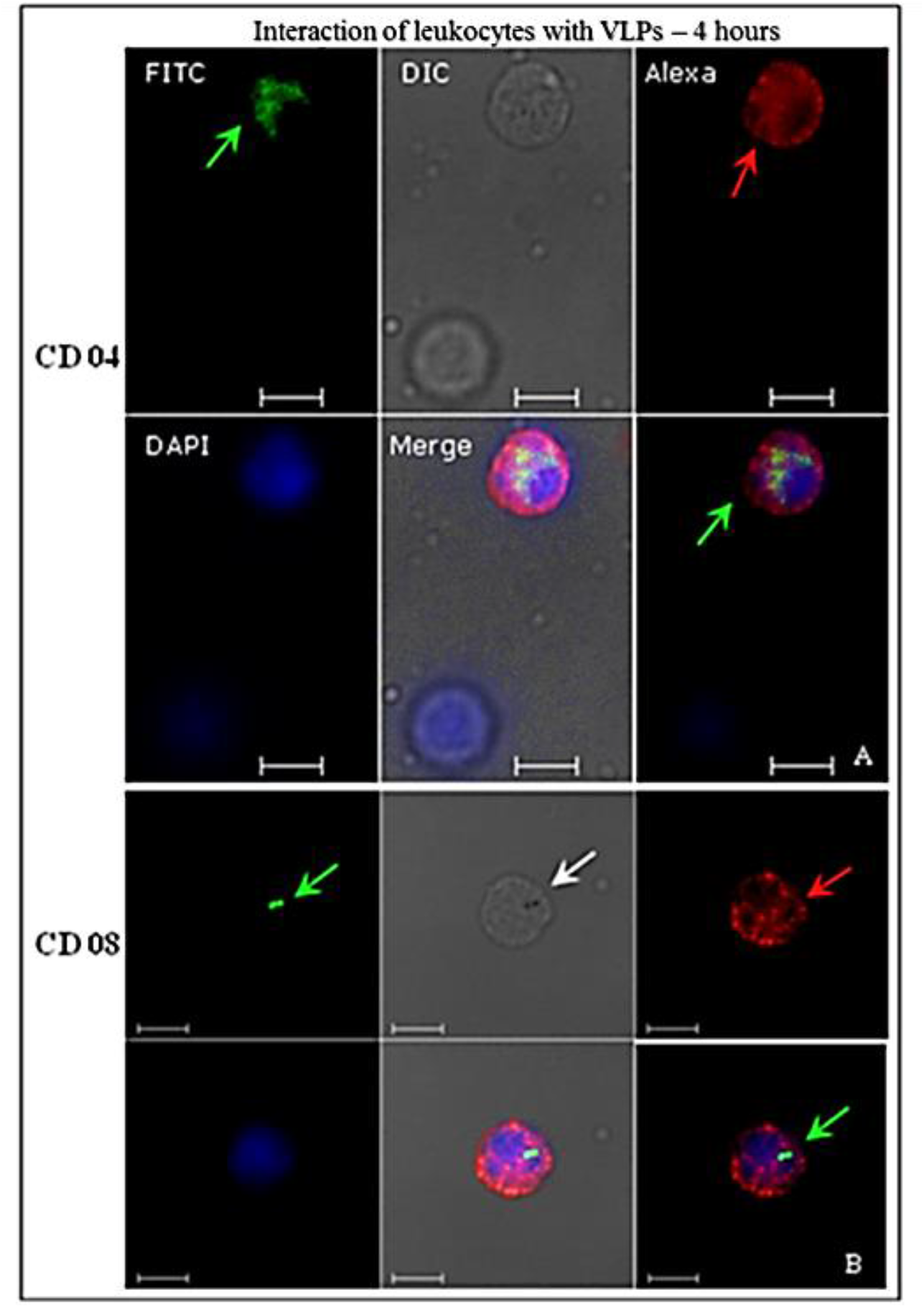
Identification of human leukocyte interactions with the HPV16 VLPs analyzed by indirect immunofluorescence. After fixation, cells were immunostained with the conformational anti-HPV16 VLPs and revealed by a secondary antibody conjugated to FITC (green). Then, the cells were treated with anti-CD04 (A) and anti-CD08 (B) and stained with a secondary antibody AlexaFluor^®^546 (red). VLP internalization (green arrows) is visible on image overlay (merged images). In (B), a structure similar to an endocytic vesicle is detectable (white arrow). CLSM Zeiss LSM 510 Meta. Magnification: Objective C-Apochromatic 63xs /1.4 oil. Bar = 5 µm.

**Figure 5:**
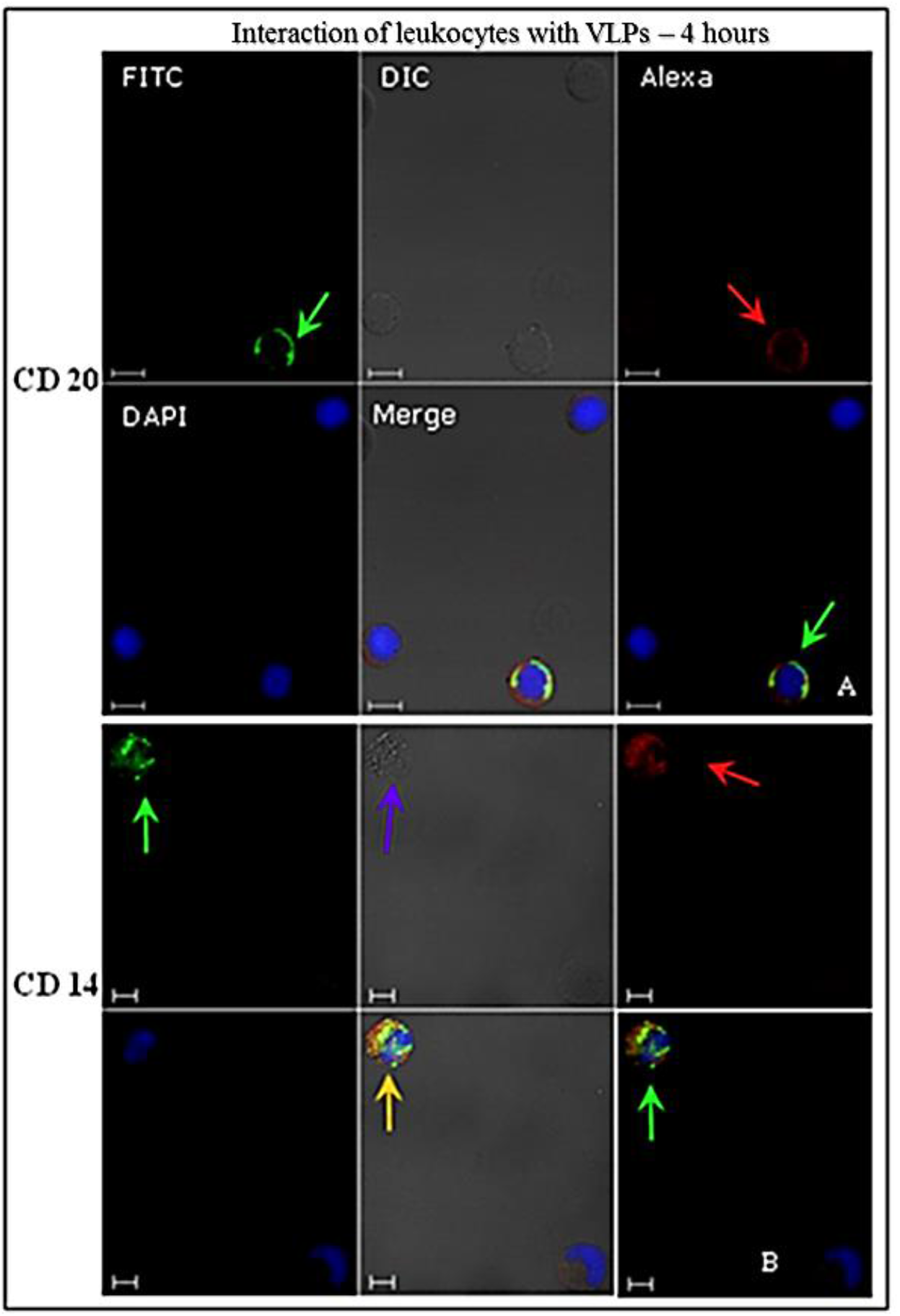
Identification of human leukocyte interactions with the HPV16 VLPs, analyzed by indirect immunofluorescence. After fixation, cells were immunostained with the conformational anti-HPV16 VLPs and revealed by a secondary antibody conjugated to FITC (green). Then, the cells were treated with anti-CD20 and anti-CD14 and stained with a secondary antibody AlexaFluor^®^546 (red). VLP internalization (green arrows) is visible on image overlay (merged images). In (B) we can observe the colocalization of CD14 and VLPs (orange arrow) and vacuole-like structures (blue arrow). CLSM Zeiss LSM 510 Meta. Magnification: Objective C-Apochromatic 63xs /1.4 oil. Bar = 5 µm.

**Figure 6:**
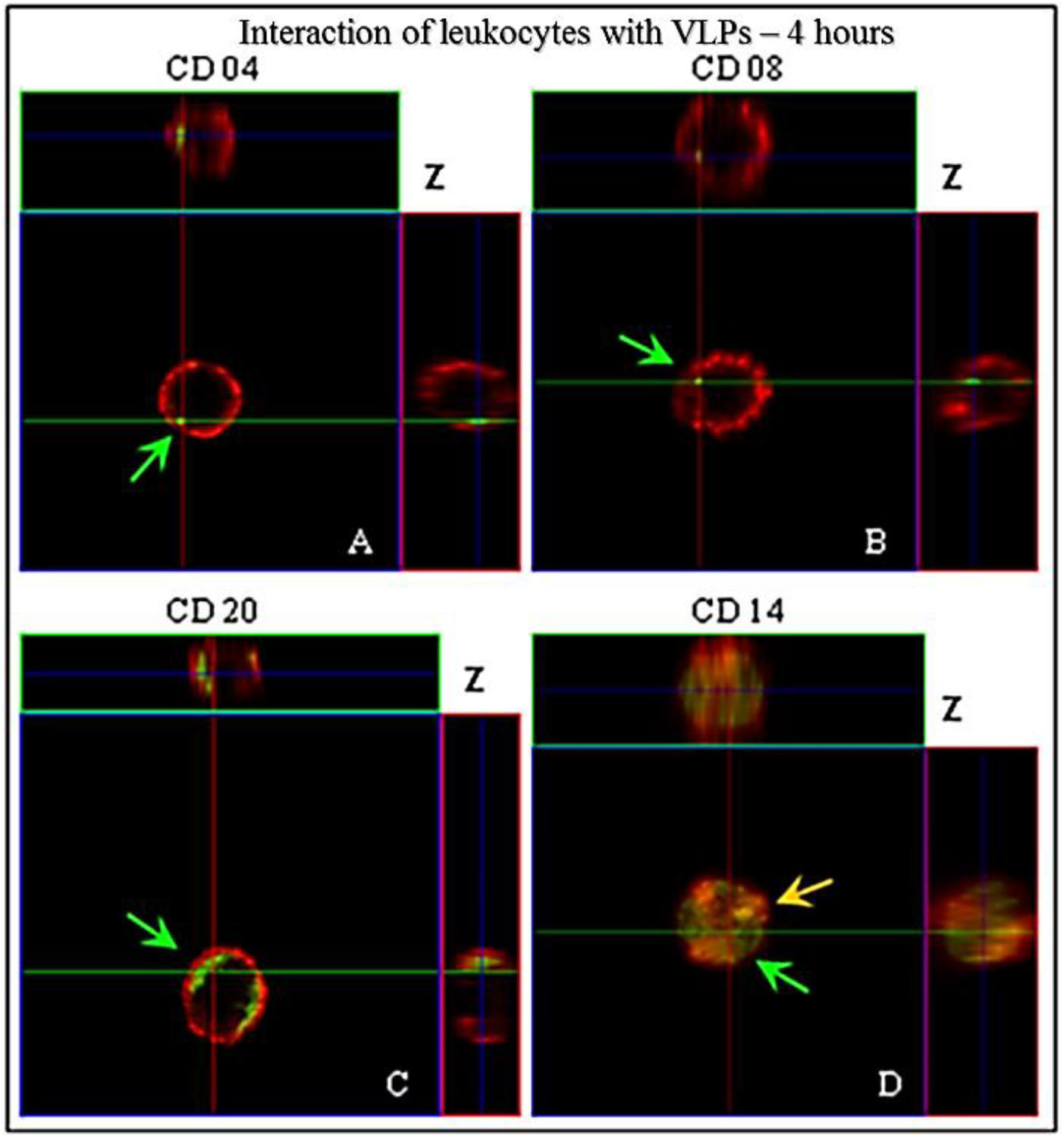
Identification of human leukocyte interactions with the HPV16 VLPs analyzed by indirect immunofluorescence. After fixation, cells were immunostained with the conformational anti-HPV16 VLPs and revealed by a secondary antibody conjugated to FITC (green). Then, the cells were treated with anti-CD20 (C), anti-CD04 (A), anti-CD08 (B), anti-CD14 (D) and AlexaFluor^®^546 revealed by secondary antibody (red). VLP internalization (green arrows) is shown in the images through Z-axis sweep cuts. (A) 9.25 µm of thick sections and each section of 0.84 µm; (B) 8.56 µm thick sections and each section with 0.95 µm; (C) 9.27 µm thick sections and each section with 1.03 µm; (D) 10,94 µm thick sections and each section having 1.40 µm. CLSM Zeiss LSM 510 Meta. Magnification: Objective C-Apochromatic 63xs /1.4 oil.

**Table 3:** Interactions of HPV16 VLPs produced in this study with PMBC (mononuclear cells) from peripheral blood of healthy female volunteers.

| PMBC | Receptor | Percentage of cells that interacted with VLPs<sup>1</sup> |
| --- | --- | --- |
| Lymphocyte T | CD04 | 52% ± 3 |
| Lymphocyte T | CD08 | 47% ± 2 |
| Lymphocyte B | CD20 | 48% ± 3 |
| Monocyte | CD14 | 23% ± 5 |

<sup>1</sup>. The percentage of cells that interact with HPV16 VLPs was calculated by the number of cells recognized by anti-CD antibodies, which have internalized the particles. The result corresponds to the analysis in duplicate, being representative of at least four trials.

### Analysis of the colocalization of VLPs and transferrin (Tf) in human PBMC

After 15 minutes, no colocalization between L1 VLPs and exogenous Tf was detected (Fig.7). Moreover, it was possible to show the internalization of VLPs during this period. After 45 minutes, colocalization of VLPs with exogenous Tf in the cytoplasm of human PBMC (Fig. 8 and 9, yellow arrows), and also with the TfR or CD71 was observed (Fig. 9 A-B, yellow arrows), suggesting that the iron pathway may be used to HPV internalization. All results were confirmed by the images overlap, respectively (Figs. 7–9).

**Figure 7:**
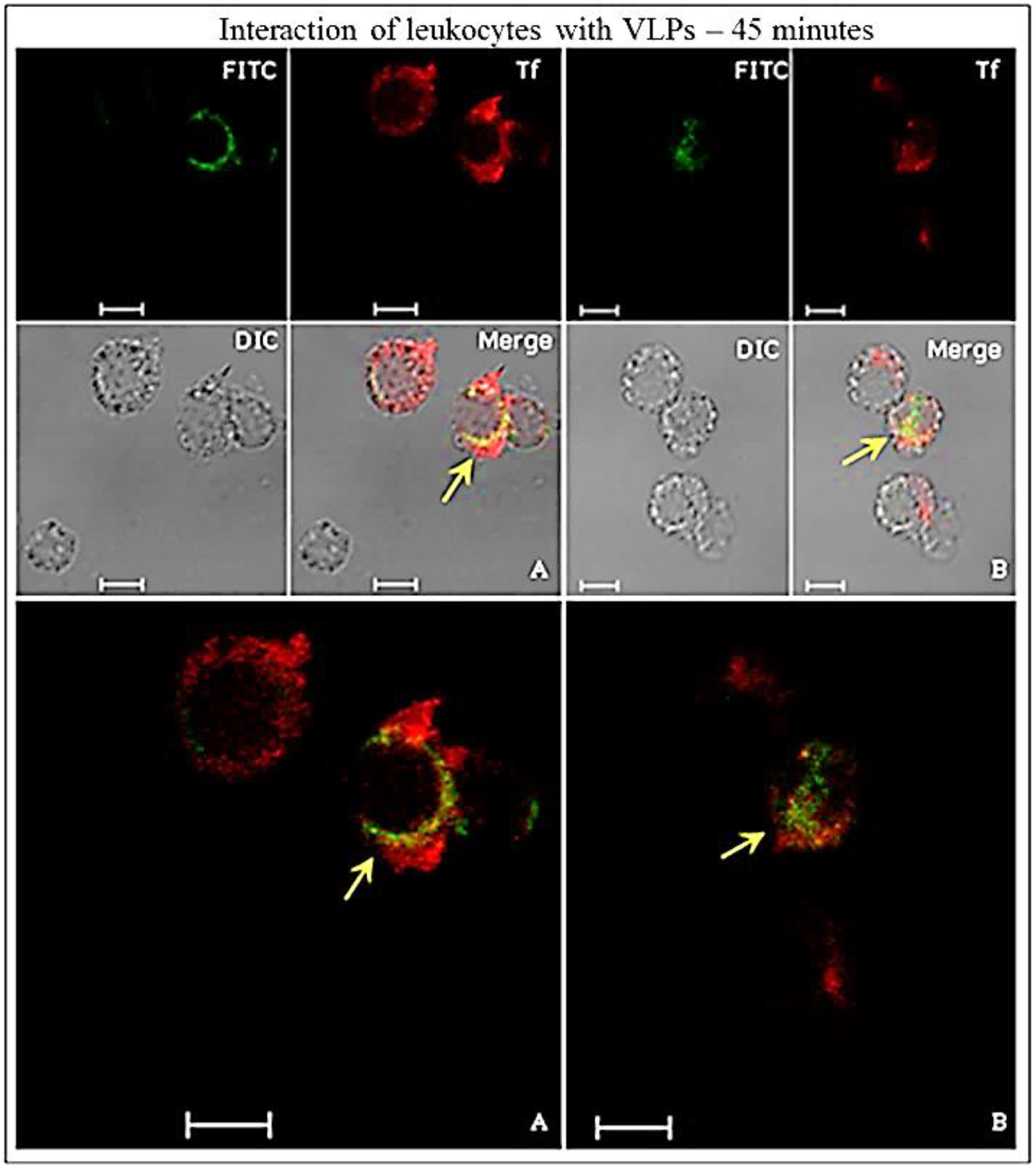
Colocalization of VLPs of HPV16 and Tf in human leukocytes, detected by indirect immunofluorescence. The leukocyte cells were incubated with the VLPs and the conjugated Tf TexasRed^®^ (red). Leukocytes were treated with anti-HPV16 L1 antibody, revealed by a secondary antibody conjugated to FITC (green). In confocality (merge), the images reveal colocalization of L1 and Tf (yellow arrows). CLSM Zeiss LSM 510 Meta. Magnification: Objective C-Apochromatic 63xs /1.4 oil. Bar = 5 µm.

**Figure 8:**
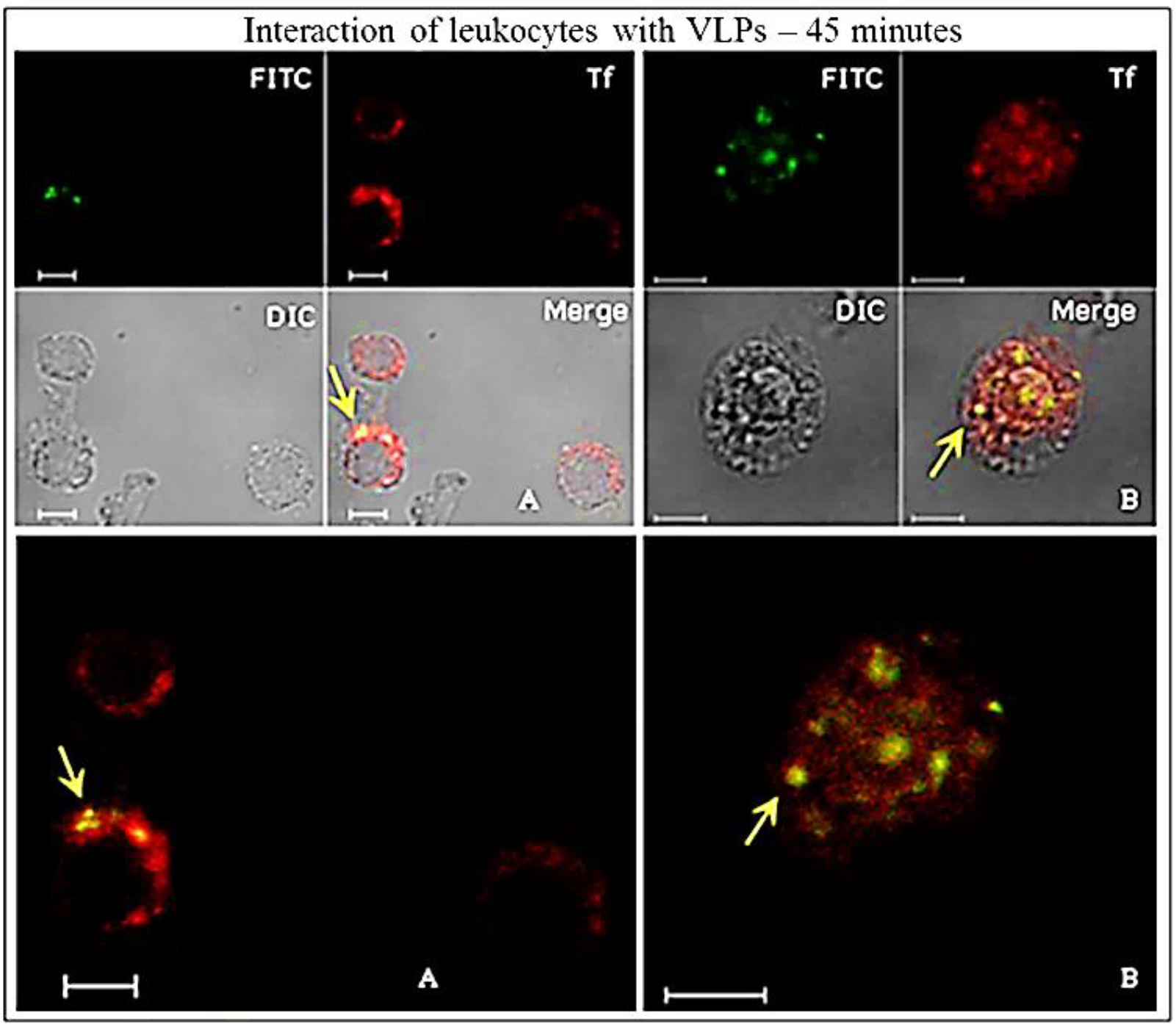
Colocalization of HPV16 VLPs and Tf in human white blood cells, detected by indirect immunofluorescence. The leukocyte cells were incubated with the VLPs and the conjugated Tf TexasRed^®^ (red). Leukocytes were immunostained with anti-HPV16 VLP antibody and revealed by a secondary antibody conjugated to FITC (green). In confocality (merge), the images reveal colocalization of VLPs and Tf (yellow arrows). CLSM Zeiss LSM 510 Meta. Magnification: Objective C-Apochromatic 63xs /1.4 oil. Bar = 5 µm.

**Figure 9:**
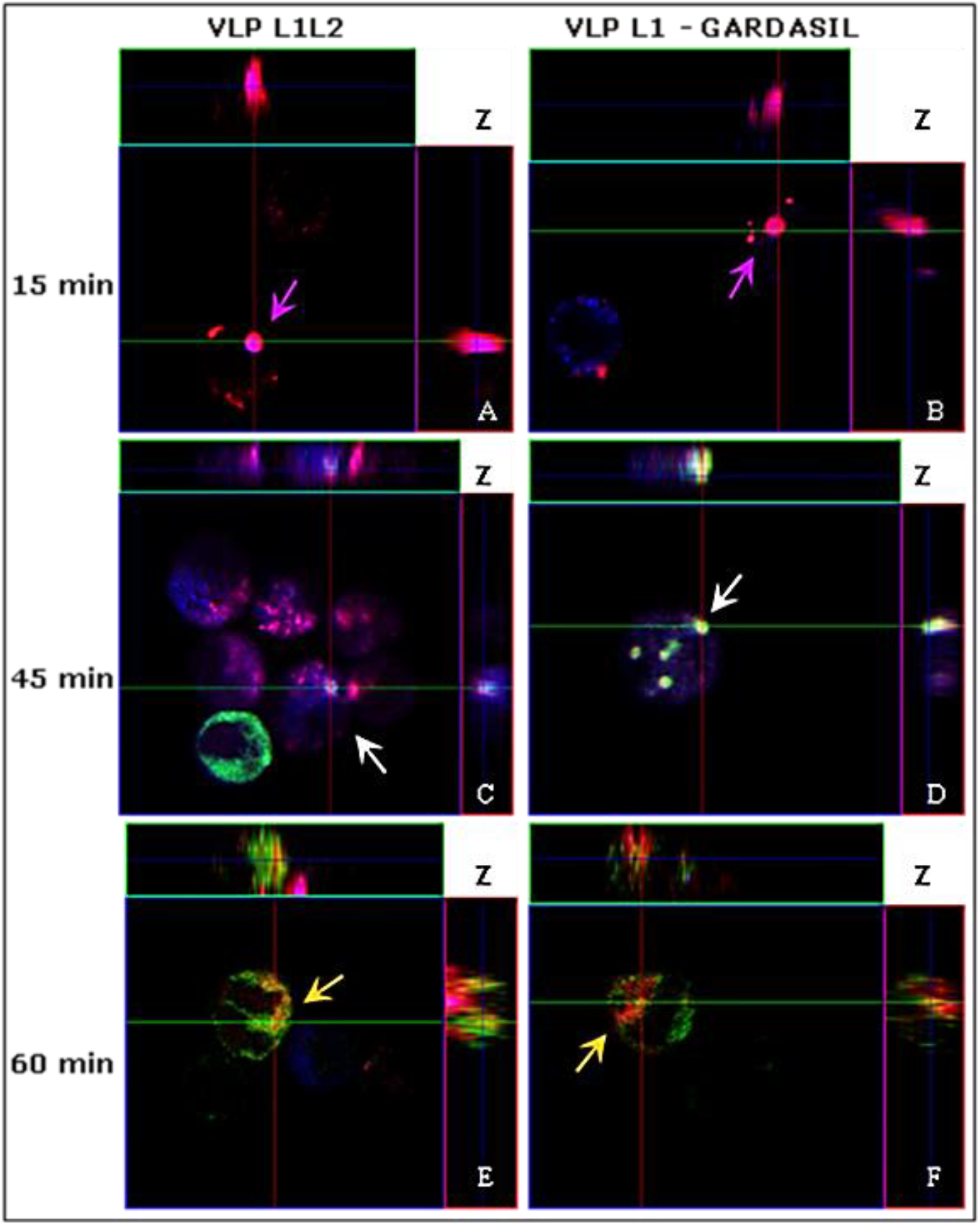
Three-dimensional imaging of leukocyte interactions with the VLPs produced in this study or the VLP vaccine Gardasil, Tf and TfR; detected by indirect immunofluorescence. The leukocyte cells were incubated with the VLPs and the conjugated Tf TexasRed^®^ (red). After fixation, the leukocytes were immunostained with anti-HPV16 VLP antibody and revealed by a secondary antibody, conjugated to FITC (green), then treated with anti-CD71 and revealed by AlexaFluor^®^633 (blue). (A) and (B) colocalization Tf and TfR (purple arrow), (C) and (D) colocalization VLP / Tf / TfR (white arrows), (E) and (F) weak colocalization VLP / Tf (orange arrows). CLSM Zeiss LSM 510 Meta. Magnification: Objective C-Apochromatic 63xs /1.4 oil.

We compared the kinetics of Tf and TfR (CD71) internalization with the colocalization of HPV16 L1/L2 VLPs produced in this study and with control HPV6, 11, 16, 18 L1 VLPs derived from the Gardasil^®^ vaccine (Fig. 9). After only 15 minutes kept in contact it was possible to observe colocalization of TfR and Tf (Fig. 9 A-B, purple arrows) with both L1/L2 VLPs (Fig. 9 A) as well as with Gardasil L1 VLPs (Fig. 9 B). By means of morphological analysis, we were able to demonstrate the colocalization of both VLPs L1 and L1/L2, with Tf and TfR in the leukocytes cytoplasm, after a 45-minutes period kept in contact (Fig. 9 C-D, white arrows). This colocalization among VLPs, Tf and TfR was found in approximately 50% of analyzed cells. Within 60 minutes in contact, we observed a feeble colocalization between Tf and VLPs in about 30% of the cells (Fig. 9 E-F, orange arrows). No colocalization was observed in the tests after 120 minutes. The colocalization was evidentiated in detail by three-dimensional figures stacked in Z-axis (Fig. 9). The negative controls showed no signs of L1 and VLPs. However, it was possible to observe exogenous Tf internalization. Comparative testing of L1/L2 VLPs produced in this study and of control L1 VLPs from Gardasil^®^ vaccine showed no significant differences in the analyzed results.

### Analysis of the blocking assays of PBMC membrane receptors

After successful identification of VLPs within leukocytes and their colocalization with Tf and TfR, a variety of biochemical inhibitors (Table 4), known to inhibit distinct cellular processes, was used to demonstrate the involvement of these and other different pathways of VLPs entry in PBMC.

**Table 4:**
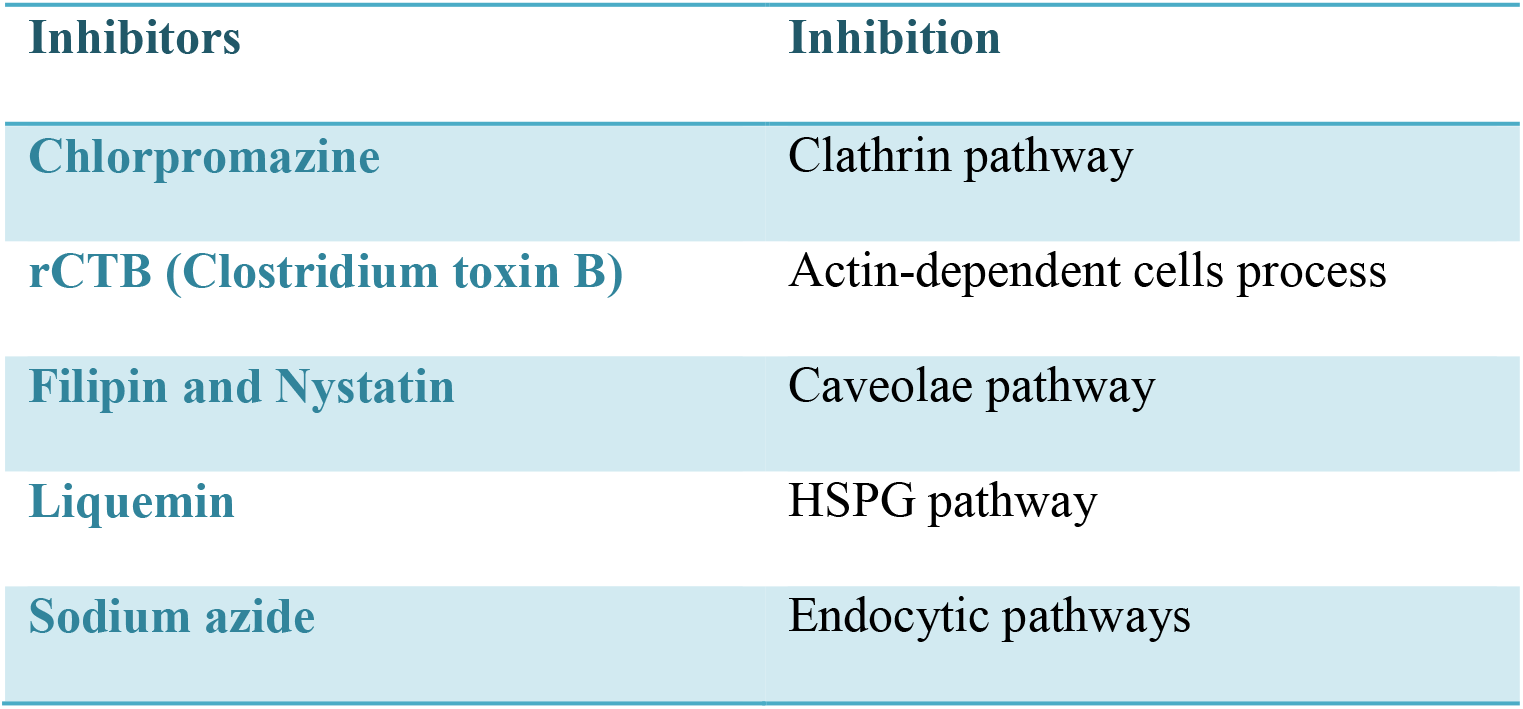
Biochemical inhibitors used to examine their effects on the entry of HPV VLPs in PBMC.

| Inhibitors | Inhibition |
| --- | --- |
| Chlorpromazine | Clathrin pathway |
| rCTB (Clostridium toxin B) | Actin-dependent cells process |
| Filipin and Nystatin | Caveolae pathway |
| Liquemin | HSPG pathway |
| Sodium azide | Endocytic pathways |

Chlorpromazine inhibits clathrin-mediated endocytosis of various plasma membrane proteins. Nystatin is a sterol-binding agent that disassembles caveolae in the membrane. rCTB (Clostridium toxin B) acts by shortening the actin filaments length, by inhibiting its process *in vitro*. Liquemin is related with the VLPs obstruction of internalization through the HSPG (Heparan sulphate proteoglycans) pathway. Sodium azide is an ATPase inhibitor, which blocks particle movement towards the cell body and leads to a diffuse random movement.

The use of these different biochemical inhibitors was not sufficient to block the HPV16 L1/L2 VLPs entry in the PBMC (Figs. 10–11). Chlorpromazine, although known for blocking the clathrin pathway through CD71 receptor of Tf, was not able to inhibit VLPs entrance inside PBMC in a satisfactory manner. The same occurred when rCTB, Filipin and Nystatin, Liquemin and Sodium azide were used to block actin-dependent cell processes, the caveolae pathway, the HSPG pathway, and endocytic pathways, respectively, either assayed separately or associated in an all-inhibitors cocktail. These results suggest that PBMC makes use of multiple internalization pathways to uptake HPV16 L1/L2 VLPs.

**Figure 10:**
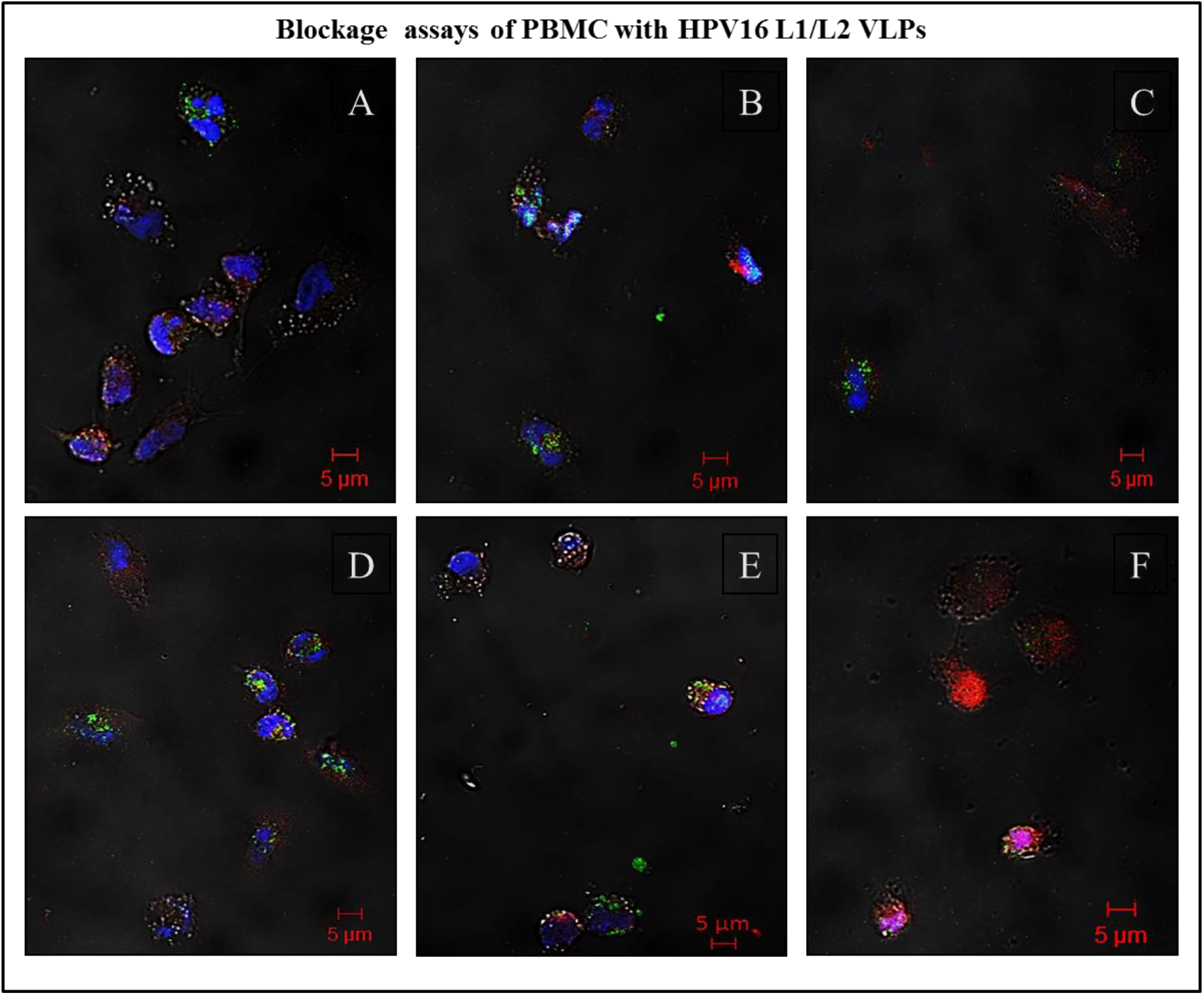
Immunofluorescence assay detecting the HPV VLPs entry in PBMC, even in presence of specific biochemical inhibitors of ligand uptake: (A) Sodium azide; (B) Chlorpromazine; (C) Liquemin; (D) rCTB; (E) Filipin; (F) Nystatin showing the overlay (merge) of images detected by stain of L1 protein (green), L2 protein (red), nuclei (blue). The entry of L1 and L2 proteins in cells was not inhibited. CLSM Zeiss LSM 510 Meta. Magnification: Objective C-Apochromatic 63xs /1.4 oil. Bar = 5 µm.

**Figure 11.**
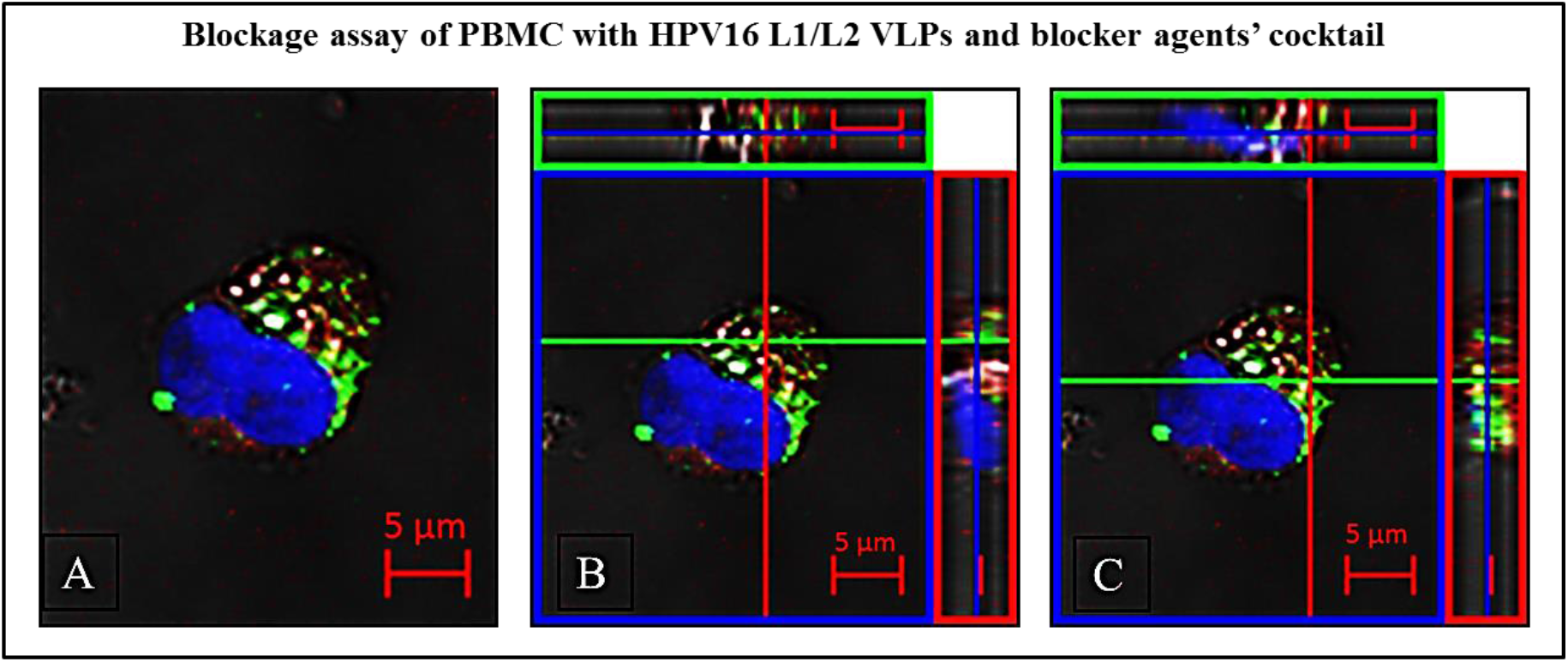
Assay performed with all inhibitors used in Fig.10 (A-F), in association. (A) Detection of L1 protein (green); L2 protein within cells (red) and nuclei (blue) in the overlay (merge) of images; (B-C) Z-axis showing the L1 (B) and L2 (C) proteins within cells. CLSM Zeiss LSM 510 Meta. Magnification: Objective C-Apochromatic 63xs /1.4 oil. Bar = 5 µm.

## DISCUSSION

Differently of many Beta and Gamma papillomavirus, Alpha papillomaviruses developed immune system evasion strategies from the host, causing persistent and visible squamous papillomas, which sometimes evolving to cancers (21–23). Some HPV types including 16, 18, 31, 33, 35, 39, 45, 51, 52, 56, 58 and 59 have been classified as human carcinogens by the IARC, being responsible for at least 4% of all human malignancies (4, 23, 24 Papillomavirus Episteme (PaVE); http://pave.niaid.nih.gov/). Infection by high-risk HPV types is the major factor for the development of cervical cancer, and HPV DNA is also found in tumors affecting other anogenital regions. Besides, they are being correlated with an increasing proportion of oropharyngeal squamous cells carcinomas, and occurrences affecting mainly tonsil and base of tongue regions are currently being documented in particular geographic areas (25–27). Cell-virus interaction is the initial step in viral infection. Often, virus adsorption occurs on the plasma membrane of susceptible cells. In general, the most common solution is that the pathogen intracellular trafficking occurs through and existing entry mechanism, mediated via clathrin, caveolin, macropinocytosis, among others, moving inside cells to reach the location where viral replication occurs (28). Transcytosis is used to move antigens and protective antibodies across epithelial barriers. Similar to what seems to occur with HIV (29), the transcytosis of HPV through epithelial cells, crossing cellular barriers, should also be dependent on trafficking to the endocytic recycling pathway. The productive life cycle of HPV is directly linked to epithelial differentiation. It is believed that the papillomavirus is capable of maintaining the expression of structural genes, under tight control of the transcription and translation regulatory mechanisms (30). The HPV L1 capsid protein, regardless of the presence or absence of L2, has the property of forming structures that mimic morphologically authentic virions, VLPs, which are being used as replacements for HPV infected cell culture *in vitro* (9, 15, 31, 32). However, the presence of L2 seems to confer greater stability to the virus and VLP, therefore, the viral capsid indeed seems to contribute to the virus infectivity (33). Most experimental models explore the process of host cell interaction by molecular or biochemical methods, using cell lines from different tissues and species in interactions with HPV L1 VLPs in their studies.

In the present work, we produced HPV L1/L2 VLPs (15, 16) to investigate the possibility that these particles to be internalized by human leukocytes. We have developed a method for the isolation of these cells (18), used in suspension at 37°C, thereby creating an *ex vivo* system as closely as possible to the natural conditions of human body, thus conceiving a novel experimental model.

HPV16 recombinant proteins L1, L2 and VLPs were visualized in the perinuclear region of HEK293T cells, in compliance with data described in the literature (34). The presence of BPV (bovine papillomavirus) in lymphocytes has been discussed (35, 36). Using PCR technique (Polymerase Chain Reaction) in samples taken from humans with anogenital lesions, the presence of HPV16 in peripheral blood samples (37) and in plasma cells (38) was detected. Besides, DNA of other HPV types has been found in PBMC (39, 40). The presence of HPV in lymphocytes of patients treated for anogenital cancers suggests that the virus can remain in these cells even after treatment. However, there is no convincing evidence supporting that lymphocytes carry/produce infectious HPV viral particles. Nonetheless, the presence of HPV macromolecules in lymphocytes may play a role in viral immune response.

The scientific literature discloses that papillomavirus exhibits specific tropism for species and tissues. However, BPV1 particles isolated from bovine warts, and HPV16 VLPs interacted successfully with 14 cell lines derived from different species and tissues, as demonstrated in two different studies (32, 41). Using our model to investigate this, HPV16 L1/L2 VLPs internalized *in vitro* and *ex vivo* with two very distinct human cells, epithelial kidney cells and PBMC, respectively. After 4-hours of interaction, these particles internalization occurred, showing a very similar distribution pattern on both cell types. In both, L1 VLPs were located in the cytoplasm near the nucleus, suggesting that tissue differences do not affect the ability of HPV16 VLPs to entry in different cell types, and that pentameric and monomeric forms of L1 may also be internalized. The interaction between VLPs and monocytes observed in this study showed colocalization with CD14. As macrophage precursors, apparently these cells are able to internalize particles at a much larger amount if compared with lymphocytes. The unexpected colocalization of CD14 membrane receptor and VLPs, as well as the morphological changes observed, suggest that monocytes respond to this interaction as immune cells, in order to eliminate invading microorganisms. CD14 is a 53-kDa molecule, expressed mainly in monocytes, macrophages and granulocytes. It is also found as soluble protein in the serum (42).

Tf is a beta globulin responsible for the transport of iron (Fe) through plasma to the interior of cells. Besides hepatocytes, other cells in the body are able to synthesize it. The TfR is a membrane protein expressed primarily in T and B lymphocytes, macrophages, and proliferating cells such as homodimers with N-terminal cytoplasmic tails (42). TfR is constantly being endocytosed by plasma membrane through clathrin-mediated endocytosis, with the purpose of carrying ferric iron bound to Tf to the inside of cells. This Tf-TfR complex is delivered to the early endosome into a lower pH medium, where iron is released and TfR is recycled to the cell surface (43). When evaluating the interactions between HPV16 L1/L2 VLPs, HPV6, 11, 16, 18 L1 VLPs (Gardasil^®^ vaccine) and unstructured HPV16 L1 in PBMCs and in the presence of exogenous Tf, we determined the colocalization of L1-VLP-Tf and free Tf in these cells, as well as colocalization with TfR, after 45 minutes of contact. This colocalization between VLP, Tf and TfR was found in approximately 50% of analyzed cells. These results suggest that HPV16 L1/L2 VLPs, HPV6, 11, 16, 18 L1 VLPs (Gardasil^®^ vaccine) and unstructured HPV16 L1 can enter cells by endocytosis through clathrin-dependent, via TfR. It is probable that, along viral infection processes, interactions between HPV16 (about 55 nm size) and Tf molecules (80-kDa) might indeed happen, through fusion with Fe-Tf-TfR complexes, mimicking its entry into the cell via low pH endosomes, thus enabling transcytosis (28, 44–47).

There have been no reports in the literature showing the internalization of HPV VLPs in human PBMC. So far, this study has showed that PBMCs of healthy female volunteers were able to internalize HPV16 VLPs. Also, T and B lymphocytes were the main cell types that further internalized VLPs. It is important to emphasize that the lymphocytes are cells capable of dividing themselves and it is estimated that the half-life of these inactive cells in humans is of some years. Furthermore, inactive lymphocytes circulate continuously through the bloodstream and lymphatic vessels. The entry of HPV VLPs in leukocytes by independent endocytosis pathways may be due to their ability to recognize, capture and remove foreign substances from the body. The use of different biochemical inhibitors, isolated or associated, does not seem to be enough to block the internalization of HPV16 L1/L2 VLPs in the PBMC. However, in order to understand the possible mechanisms involved in these PBMC interactions in future studies, which in turn cannot be assessed only by VLP experiments, it will be necessary to drawn new strategies using PsV (pseudovirion) infectious particles provided with HPV16 viral genome (48–50). Thinking in another approach, it will be possible to exploit this experimental model in vaccines-cells interaction studies and the immunological mechanisms involved, for instance, evaluating the impact of vaccines and the prevalence of determined viral genotypes in distinct populations (51–54). A recent review discusses interestingly the importance to clarify some lapses of our understanding about HPV natural history (55).

## CONCLUSIONS

HPV16 L1/L2 VLPs produced in this study interacted with *in vitro* HEK293T cells and *ex vivo* PBMC from healthy women volunteers, and were internalized. This study evinced that HPV16 L1/L2 VLPs can interact with the plasma membrane surface, been uptake by lymphocytes. We also demonstrated that HPV16 L1/L2 VLPs might utilize the endocytic pathway of iron-mediated clathrin as a receptor, which involves multiple membrane receptors simultaneously and also the classical endocytic pathway of iron, the most widespread among vertebrates and the animal kingdom as a whole, we have demonstrated that HPV16 L1/L2 VLPs does not require a specific receptor in order to be internalized by leukocytes. Complementary studies are required to accurately demonstrate the probable mechanisms involved.

## AUTHORSHIP

A.M.C. conceptualized and wrote the manuscript.

## ACKNOWLEDGEMENTS

The authors would like to express their deepest gratitude to the CAPES Master Degree Fellowship to V. Szulczewski and E.A. Kavati, from the Biotechnology Interunits Postgraduate Program USP-IBU-IPT; PAP-SES-FUNDAP Fellowships to T.M. Hosoda; Financial Support: FAPESP – Process Number 2004/15122-5; Butantan Foundation and Butantan Institute; to the Prof Dr. Richard B.S. Roden, from the Department of Pathology, Johns Hopkins University, Baltimore, MD, USA, which kindly provided us anti-L2 antibody, and for useful comments on the manuscript.

## ETHICAL COMMITTEE

This work was duly evaluated and approved by the Sao Joaquim Hospital, Real and Meritorious Portuguese Beneficence Association Ethical Committee on Human Research. Protocol number: 369-08.

This work is in full compliance with the National Biosafety Law (CTNBio) guidelines and the present study was approved by the Butantan Institute Institutional Biosafety Committee (CIBio) and CTNBio – Process Number 01200.004893/1997-93, published in the Brazilian Official Gazette (DOU) on August 16, 2011.

## ABBREVIATIONS

HPV: human papillomavirus
VLP: virus-like particles
L1: major capsid protein
L2: minor capsid protein
HEK293T: human epithelial cell lineage 293T
CD71: transferrin receptor
CD4: T lymphocyte membrane receptor
CD8: T lymphocyte membrane receptor
CD20: B lymphocyte membrane receptor
CD14: monocyte membrane receptor
rCTB: recombinant cholera toxin B subunit is the non-toxic portion of cholera toxin (CT)
HIV: human immunodeficiency virus
pUF3L1h vector: humanized expression vector for L1 protein
pUF3L2h vector: humanized expression vector for L2 protein
EDTA: anticoagulant
Tf: transferrin
DMEM: Dulbecco’s Modified Medium
FBS: fetal bovine serum
RPMI: Roswell Park Memorial Institute cell culture medium
PBMC: peripheral blood mononuclear cells
BSA: bovine serum albumin
TEM: transmission electron microscopy

